# Taxon disappearance from microbiome analysis indicates need for mock communities as a standard in every sequencing run

**DOI:** 10.1101/206219

**Authors:** Yi-Chun Yeh, David M. Needham, Ella T. Sieradzki, Jed A. Fuhrman

## Abstract

Mock communities have been used in microbiome method development to help estimate biases introduced in PCR amplification, sequencing, and to optimize pipeline outputs. Nevertheless, the necessity of routine mock community analysis beyond initial method development is rarely, if ever, considered. Here we report that our routine use of mock communities as internal standards allowed us to discover highly aberrant and strong biases in the relative proportions of multiple taxa in a single Illumina HiSeqPE250 run. In this run, an important archaeal taxon virtually disappeared from all samples, and other mock community taxa showed >2-fold high or low abundance, whereas a rerun of those identical amplicons (from the same reaction tubes) on a different date yielded “normal” results. Although obvious from the strange mock community results, due to natural variation of microbiomes at our site, we easily could have missed the problem had we not used the mock communities. The “normal” results were validated over 4 MiSeqPE300 runs and 3 HiSeqPE250 runs, and run-to-run variation was usually low (Bray-Curtis distance was 0.12±0.04). While validating these “normal” results, we also discovered some mock microbial taxa had relatively modest, but consistent, differences between sequencing platforms. We suggest that using mock communities in every sequencing run is essential to distinguish potentially serious aberrations from natural variations. Such mock communities should have more than just a few members and ideally at least partly represent the samples being analyzed, to detect problems that show up only in some taxa, as we observed.

**Importance:** Despite the routine use of standards and blanks in virtually all chemical or physical assays and most biological studies (a kind of “control”), microbiome analysis has traditionally lacked such standards. Here we show that unexpected problems of unknown origin can occur in such sequencing runs, and yield completely incorrect results that would not necessarily be detected without the use of standards. Assuming that the microbiome sequencing analysis works properly every time risks serious errors that can be avoided by the use of suitable mock communities.

## Introduction

Analysis of microbial community composition by 16S or 18S rRNA tag sequencing is well recognized as a powerful tool to evaluate microbial diversity in virtually all environments (1–6). By taking advantage of high-throughput sequencing, microbial ecologists can now easily reveal microbiomes with high coverage, including the “rare biosphere”. Some potential problems are well recognized, such as chimeric sequences derived from PCR amplification artifacts (7, 8) and random sequencing errors (9, 10). A number of studies thus have developed various pipelines and algorithms to detect chimeras and other likely errors, and remove them from downstream analyses (9–14). Such processing strategies help to correct or remove some errors within each sequencing run, yet they do not verify if each sequencing and amplification analysis consistently works the same way each time and thus retrieves comparable measures of the community composition. As many microbiome projects and meta-analysis studies process sequences from multiple sequencing runs, consistency between analytical runs is critical for properly compiling data from different labs and different runs. Otherwise, the errors may be propagated to future studies.

In general, biases and many kinds of potential analytical errors can be detected by preparing and sequencing mock communities, which can act as known standards to be used during sample analysis. This is analogous to a set of standards of the sort used in virtually every careful chemical or physical assay, for instrument “calibration” and/or to test that a particular batch or reagents is working as expected. Such mock communities can be genomic preparations (15) or collections of known 16S rDNA gene fragments or clones (16), which may be used for a variety of purposes, but in general represent known standards. Since the abundance of OTUs in mock communities is known *a priori*, such mock communities can be used initially to test for biases during method development, and also can be used in each run to verify that the analysis is within acceptable bias limits (16).

Although mock communities have been used to characterize biases in community analyses (15, 16) and support use of highly-resolving analysis approaches (17), they are still not commonly involved in empirical studies. Why such standards are not routine in microbiome analysis is not fully clear; often it seems to be assumed that careful replication of methods, such as using the same DNA extraction, PCR amplification and sequencing methods throughout a given study would yield sequences merely containing random minor errors that are consistent between samples. Additionally, while experimental procedures and collection are usually performed by the lab doing the study, library preparation and/or sequencing is often performed off-site at an academic or commercial sequencing facility, and to some extent in blind-faith. But today with the low cost of sample preparation and analysis for sequencing, there would seem to be few excuses not to use such standards. Contrary to the expectation that the sequencers work the same every time, in this study we found a remarkably aberrant sequencing result, showing that an important marine taxon almost disappeared in mock communities and field samples from one sequencing run but was recovered in another run using the exact same PCR products. Suitable diverse mock communities offer a good chance to detect such errors and to help validate each batch of results.

## Results and Discussion

We have been using mock communities for over 4 years, primarily on the Illumina MiSeq platform, with generally consistent results from run to run (16, 17). However, we were surprised to notice that in one run analyzed on a HiSeqPE250 (summer 2016), the mock communities yielded a completely unexpected result whereby the Marine Group II archaea were virtually absent, and other taxa had quite unexpected relative abundances whether clustered into 99% OTUs (Fig. 1) or amplicon sequence variants (ASVs, Fig. S1). We found the same result from samples prepared by three different individuals, each of whom worked independently and from different aliquots of the same mock community materials. This led us to reanalyze the samples and to also carefully compare several mock community runs on both the MiSeq and HiSeq platforms.

**Figure 1.**
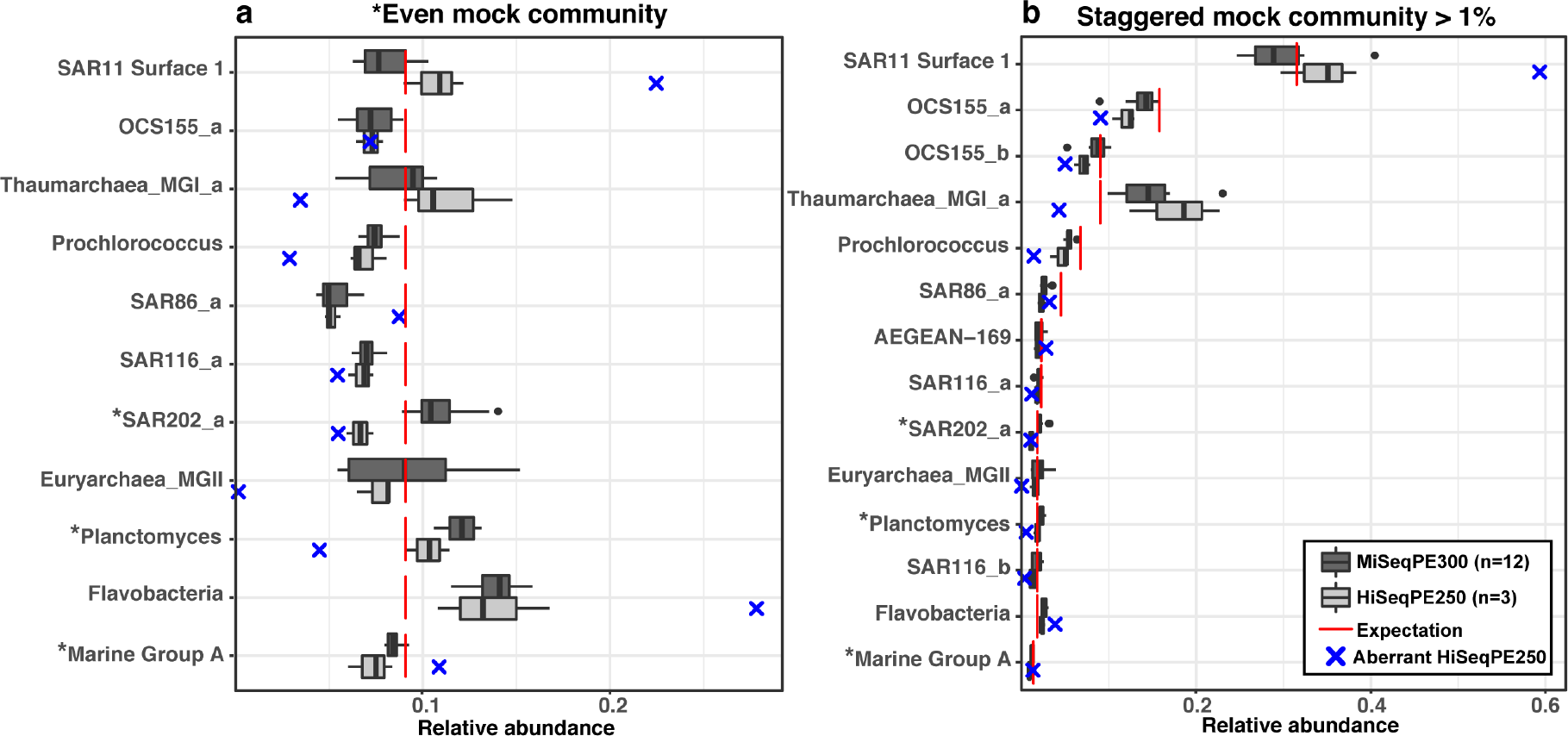
Comparisons of a) even mock communities and b) staggered mock communities sequenced by MiSeqPE300 and HiSeqPE250. Significant differences between MiSeqPE300 versus HiSeqPE250 of each clone are further marked with an asterisk on the name (Wilcoxon-rank sum test, p<0.05). Significant difference in the whole-community composition between MiSeqPE300 and HiSeqPE250 was only found in the even mock community (ANOSIM test, p<0.05)

An obvious first question to ask is whether this unusual result was simply due to the use of a HiSeq rather than MiSeq as had been used previously. Comparison of 3 additional HiSeq runs to 4 MiSeq runs (with multiple replicates in each run) showed that there was overall run-to-run consistency (Bray-Curtis distance was 0.11±0.04 in the even mock community and 0.12±0.04 in the staggered mock community), with HiSeq and MiSeq producing statistically indistinguishable whole-community outcomes in “staggered” mock communities (27 taxa at different abundances), yet there were relatively modest but statistically significant differences in “even” mock communities (11 taxa at the same abundance; ANOSIM test, p < 0.05). When examined individually, clones in both “even” and “staggered” mock communities representing SAR202, Planctomyces, and Marine Group A, as well as OCS155_b which was included only in the staggered mock communities, were significantly different between the two sequencing platforms, in at least one comparison of OTUs (Fig.1) or ASVs (Fig. S1) (Wilcoxon-rank sum test, p < 0.05). Even for the significantly different ones, the discrepancy between platforms was generally modest, with the results generally differing by less than 1.3 fold, though the SAR202 clone differed by ~1.67 fold. However, this sharply contrasted with the aberrant HiSeq sequencing run, in which mock community members representing Euryarchaea Marine Group II almost disappeared (i.e. 0 in the staggered mock community when it should have been ~1.8% and ~0.15% in the even mock community when it should have been ~9%), Thaumarchaea and Prochlorococcus underweighted more than 2-fold, and SAR11 and Flavobacteria were overweighted more than 2-fold (Fig. 1).

To test if this bias was introduced in the sequencing process step itself (rather than in sample preparation up to and including PCR), the same PCR products of the aberrant mock communities (along with PCR products of field samples which were also included in the aberrant run) were resequenced on MiSeqPE300, and we indeed found sequence abundances returned to “normal,” i.e. indistinguishable from the other MiSeq runs, whether clustered by 99% OTUs (Fig. 2) or ASVs (Fig. S2). Note that with our protocols, the PCR products themselves were “ready to run” as sequencing libraries (and simply mixed with other samples that had different barcodes), so there were no additional library preparation steps that could have altered the relative compositions.

**Figure 2.**
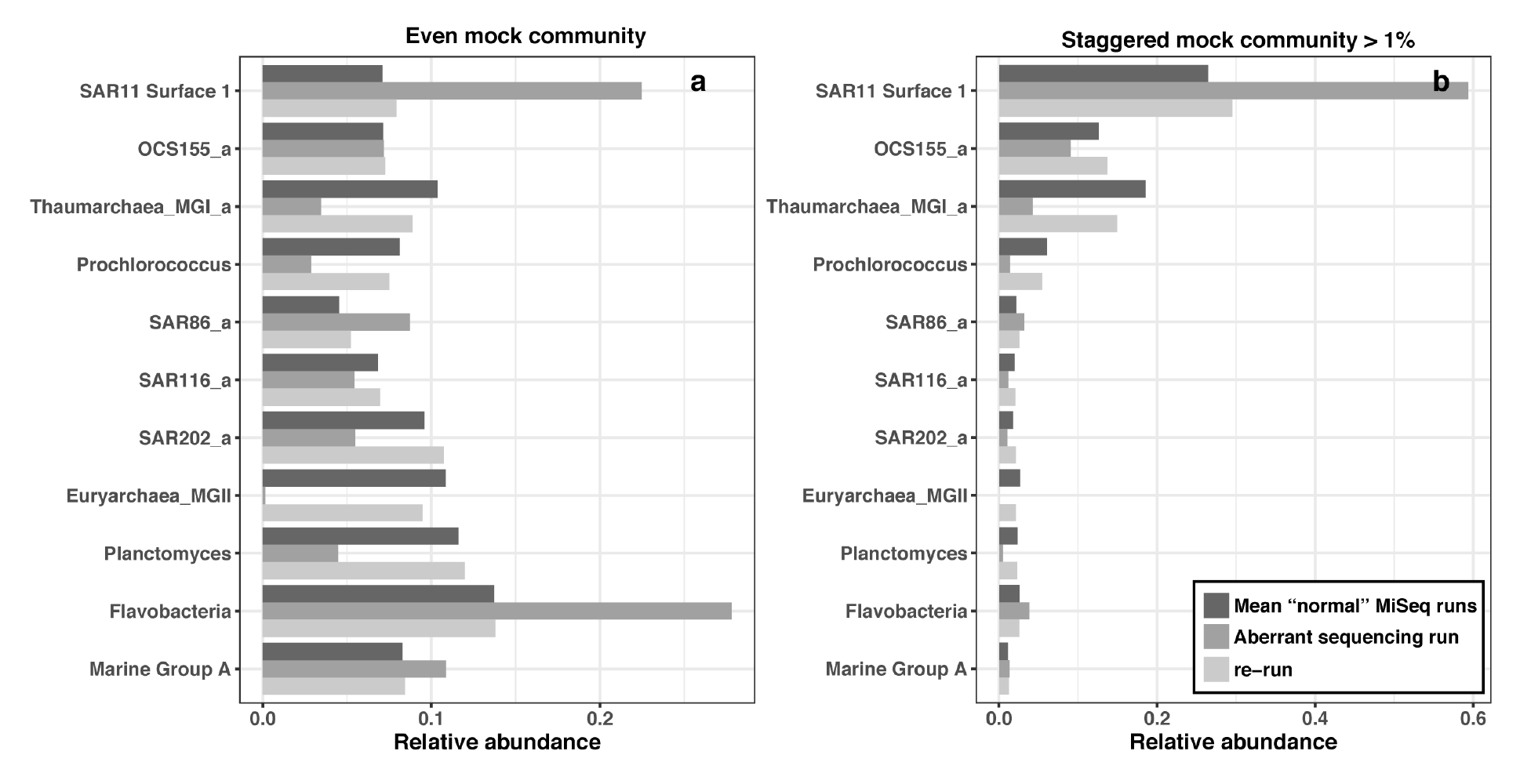
Re-run of the same PCR products from the “aberrant” sequencing run.

To test if the biases observed with the mock communities were also found in field samples, two field samples were analyzed multiple times and compared (Fig. 3). When “normal” sequence runs were compared by analyzing a field sample (e.g. a surface seawater sample collected in April 2013 at the San Pedro Ocean Time-series (SPOT) location), the results showed that the rank abundance curves were generally similar between sequencing platforms and even between slightly different primers (Bray-Curtis distance was 0.11±0.04). However, when we compared a field seawater sample collected in June 2015, which is included in the aberrant sequencing run, and re-sequenced the same PCR product (i.e. from the same original tube), the rank abundance curves were substantially different considering they are supposed to be replicates (Bray-Curtis distance was ~0.31). The top 20 abundant OTUs showed that in the aberrant run SAR11 OTUs were overweighted, and Euryarchaea Marine Group II was missing - the same pattern we found in the mock communities (Figs 1, 2, 3b). Interestingly, in the aberrant run, three different SAR11 OTUs were strongly overweighted, two different MGII Archaea were strongly underweighted, as were five different SAR116 taxa, suggesting the biases were group-specific (Fig 3b). Despite these differences which appeared clear by re-sequencing the identical sample, we note that had we not been alerted by the aberrant mock community results, the field sample results themselves did not appear so unusual overall; aside from the missing MGII archaea (which might not have been noticed, or might have thought to be real), most taxon abundances fell within the natural variation in our study area, where Bray-Curtis distances between near-surface communities typically range ~0.2-0.6 (18). Hence it was the mock community standards that revealed the problem.

**Figure 3.**
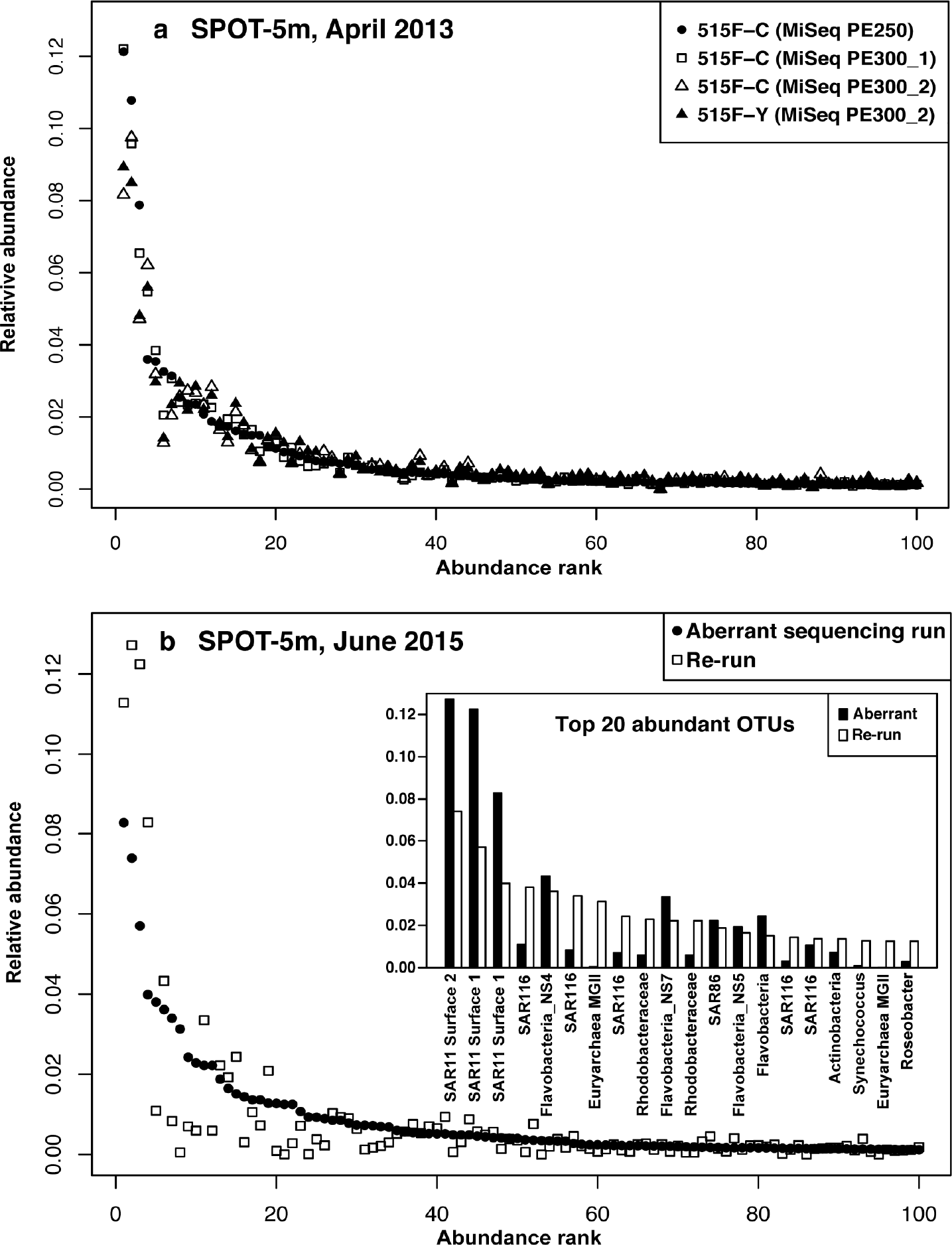
Field community comparisons via 16S and between sequencing runs. a) good replication of rank abundance curves between different sequencing runs and with slightly different primers (515F-C is the original EMP primer and 515F-Y is the version where a C is replaced with a Y, (16)). b) Comparison of rank abundance curves from the June 2015 sample analysis, showing the aberrant sequencing run and the exact same PCR products re-analyzed on the other sequencing run on a different day.

## Conclusions

Our results suggest that including mock communities as standards in every sequencing run is strongly advised, since it is the only way to verify each sequencing run is “behaving normally,” which we showed is not always the case. Ignoring that possibility may lead to serious errors that may obscure real patterns and lead to erroneous conclusions. Even duplicate sequencing in different runs would not help determine which data are “correct” when two runs are significantly different. While the “missing” archaeal taxon in our study might represent a “smoking gun” of the sort that could raise concern by researchers paying close attention to each result, there could easily be less obvious changes in a given sequencing run that could strongly bias results without being noticed. In our case, because of the aberrant mock community outcomes, we were able to objectively discard the results of a run. Mock communities can also show real, even subtle, differences between analytical protocols, as we found when comparing the MiSeq and HiSeq sequencing platforms. Because we found that only a few taxa (though common in seawater) showed the strongest biases, we suggest that mock communities should include several representative taxa to better increase the chance of detecting potential problems.

## Materials and Methods

### Sample collection and DNA extraction

Samples were collected from 5 m depth at the San Pedro Ocean Time-series (SPOT) location in April 2013 and June 2016. ~12 L of seawater was pre-filtered through an 80-μm mesh to remove metazoa and was then sequentially filtered through a 1.2 μm A/E filter (Pall, Port Washington, NY) and a 0.2 μm Durapore filter (ED Millipore, Billerica, MA). The filters were stored at −80°C until DNA extraction.

### Mock communities preparation

To generate even and staggered mock communities, 11 or 27 clones of marine 16S rDNA were prepared respectively (16), briefly as follows: Clones were originally generated from 16S-ITS-23S amplified products from marine DNA. The plasmids were purified from clones and amplified with M13F and M13R primers. Then, bacterial 16S PCR products were generally amplified with 27F and 1492R primers, and archaeal with 20F and 1392R, in order to get nearly full-length products. In the even mock community, the DNA mixture had an equal amount of each PCR product (11 in total). In the staggered mock community, the DNA mixture had different proportions of each PCR product (27 in total), roughly mimicking the marine bacterioplankton distribution from our sample site.

### PCR and sequencing

To pool multiple samples in a single Illumina paired-end sequencing platform, a dual-index sequencing strategy was used. The V4 and V5 hypervariable regions of the 16S rDNA gene were amplified using the forward primer *A-I*-NNNN-barcode-515F (*A-I*-NNNN-barcode-GTGYCAGCMGCCGCGGTAA) and reverse primer *A*-index-*I*-926R (*A*-index-*I*-CCGYCAATTYMTTTRAGTTT), where *A* is the Illumina sequencing adapter, *I* is the Illumina primer, and barcode and index are sample specific tags (5 bp barcode and 6 bp index). For each sample, one 25 μL amplification mixture contained 1.25X 5Prime Hot Master Mix (0.5 U taq, 45 mM KCl, 2.5 mM Mg^2+^, 200 μM dNTPs), 0.3 μM primers, and 0.5 ng of DNA sample. The PCR conditions were an initial denaturation at 98°C for 2 min; 30 cycles of 98°C for 45 s, 50°C for 45 s, 72°C for 90 s; and a final extension at 72°C for 5 min. Each PCR product was cleaned using 0.8X Ampure XP magnetic beads (Beckman Coulter). Purified PCR products from samples were quantified with PicoGreen and then sequenced on Illumina HiSeq 2500 in PE250 mode and/or MiSeq in PE300 mode. For each sequencing run, multiple blanks and two versions of mock communities (even and staggered) were included as internal controls.

### Sequencing output processing

Sequences were demultiplexed by reverse index allowing for one mismatch, at the sequencing facility. Then, the forward barcodes were extracted using QIIME 1.9.1 *extract_barcode.py* (19). The forward and reverse reads were de-multiplexed with forward barcodes independently allowing no mismatch using QIIME 1.9.1 *split_libraires_fastq.py*. The fully demultiplexed forward and reverse reads were then split into per-sample files using QIIME *split_sequence_file_on_sample_ids.py*. The raw sequences after being demultiplexed and split into per-sample fastq files have been submitted to the EMBL database under accession number PRJEB12267 and PRJEB22835. The demultiplexed forward and reverse reads were quality filtered using Trimmomatic 0.36 (SLIDINGWINDOW:4:20 MINLEN:200) (20) and merged using USEARCH v7 *fastq_mergepairs* (21). The forward and reverse primers were then trimmed from the merged reads using cutadapt (22). Chimeric sequences were identified and removed by *de novo* chimera checking using QIIME 1.9.1 *identify_chimeric_seqs.py* and *filter_fasta.py*. Before clustering, we added to the sequences an artificial file (*in silico expected relative abundance*) containing the mock community sequences in their exact proportion and sequence composition (a “perfect” mock community), in order to help trace the outcome of the sequenced mock communities through the clustering and OTU table generation. Operational taxonomic units (OTUs) were clustered at 99% similarity cut-off by UCLUST within QIIME 1.9.1. The most abundant sequence of each OTU was chosen as the representative sequence. The taxonomy of each OTU was assigned with reference-based UCLUST against SILVA v119 database (23) using QIIME *assign_taxonomy.py*. As an alternative to OTU clustering, we also implement minimum entropy decomposition (MED) (14) on our sequence set to generate Amplicon Sequence Variants (ASVs) that differ from each other at specific bases (as distinct from OTUs that can differ at any base). In brief, MED aims to recognize real genetic variants from sequencing error, to partition the community at fine phylogenetic resolution. In our analysis, we used 0.25 as the entropy threshold to distinguish real variances from sequences error based on our previous work at SPOT (17). We only considered ASVs that had at least 50 individuals represented across all samples examined. Analysis by OTU clustering and MED yielded the same conclusions, and we include MED results in the supplemental materials.

## Acknowledgments

We thank Erin Fichot, Alma E. Parada, and Julio C. Ignacio-Espinoza for help with sampling and laboratory work. This work was supported by NSF grant 1136818 and grant GMBF3779 from the Gordon and Betty Moore Foundation Marine Microbiology Initiative.

